# Common brain network dynamics capture attention fluctuations in tasks and movies

**DOI:** 10.1101/2025.09.18.677177

**Authors:** Anna Corriveau, Jin Ke, Monica D. Rosenberg

## Abstract

Attentional states are highly dynamic and variable, fluctuating from moment to moment and showing stark differences across contexts. To what extent does functional brain reorganization capture variability in attentional states? In the present study we utilize a time-resolved measure of functional MRI connectivity to examine and compare the extent to which univariate activity and functional networks reflect second-to-second sustained attentional fluctuations. Sustained attention was measured objectively, using auditory and visual tasks, and subjectively while participants watched and listened to narratives. Results revealed that objective measures of sustained attention to images and sounds involved common patterns of neural activity and functional interactions. Additionally, networks related to sustained attentional performance during controlled tasks also predicted fluctuations in subjective attentional engagement while participants watched movies and listened to a podcast. Generalization between experimental and everyday task contexts highlights the robustness of time-resolved functional networks for capturing dynamic fluctuations in sustained attentional states.

## Introduction

Large-scale functional brain networks are constantly in flux. One ongoing goal of cognitive neuroscience is to relate dynamic neural fluctuations to changes in internal states and behaviors. To date, identification of dynamic functional networks has been limited in its temporal resolution, requiring calculation of functional connectivity over time windows. Recent methods, however, enable the exploration of functional network reconfiguration at individual points in time, removing this constraint (Faskowitz et al., 2020; Zamani Esfahlani et al., 2020). In the current study, we exploit time-resolved network analyses to identify functional networks that track fluctuations in sustained attention, which occur over a matter of seconds (Rosenberg & Song, 2021).

While sustained attention is ubiquitous throughout daily life, situations requiring continuous focus may differ in many ways. What we attend, the perceptual modality in which information is received, and the salience of information may differ between contexts, among other factors. For example, while reading an email and conversing with a friend both require the ability to attend and understand a stream of words, these activities require processing of different perceptual modalities and may differ dramatically in their inherent ability to exogenously capture attention. Despite these discrepancies, work examining individual differences shows that sustained attention performance is consistent across task contexts (Unsworth et al., 2021) and perceptual modalities (Corriveau et al., 2025b; Seli et al., 2011; Terashima et al., 2021), supporting a core sustained attentional construct. Therefore, generalizable brain-based models of sustained attention must capture context- and modality-agnostic features of sustained attention.

Interactions between brain regions measured using functional magnetic resonance imaging (fMRI) have repeatedly been linked to sustained attention. Functional MRI connectivity-based networks capture differences in sustained attention ability across individuals (Corriveau et al., 2022; Rosenberg et al., 2016) and are highly generalizable, predicting individual differences across different attention tasks (Yoo et al., 2022), perceptual modalities (Corriveau et al., 2025a), and age groups (Kardan et al., 2022). However, sustained attention not only varies across individuals but also within an individual on the order of seconds (Castellanos et al.., 2005; Rosenberg et al., 2025; Terashima et al., 2021). Work investigating fluctuations over time using block-averaged connectivity (Rosenberg et al., 2020; Kardan et al., 2022), sliding-window connectivity (Song et al., 2021), or psychophysiological interaction analysis (Kucyi et al., 2017) has shown that functional connections reconfigure with changes in attentional state. These previous methods, however, involve averaging or smoothing across time windows which may obscure high-frequency fluctuations in attention.

A recently-developed method, termed edge time series, computes functional interactions between brain regions at a single time point, eliminating the need to average or smooth functional connectivity data over time (Faskowitz et al., 2020; Zamani Esfahlani et al., 2020). In fMRI, edge time series reflect the dot product of z-scored univariate activity time courses. Because this calculation can be performed on a single time point, edge time series enable the identification of functional connections that correspond with high-frequency changes in cognition and behavior. Using methodology akin to univariate general linear models (GLMs), edge-based GLMs can identify networks of edges whose interaction, or co-fluctuation, tracks differences in sustained attentional state from trial-to-trial (Jones et al., 2024). Importantly, while mathematically equivalent to the interaction between univariate time courses, edge time series provide explanatory power above and beyond univariate activity (Merritt et al, 2024). Therefore, edge time series enable the identification of neural substrates of dynamic cognition and attention at a higher resolution than previously thought possible.

An additional barrier to the identification of functional networks related to cognition and attention is their generalization. Valid functional networks should capture variation in a new dataset that may differ in many ways from data used to train the networks. With the recent development of edge-based GLMs, relatively little work has tested their ability to generalize across task contexts. Here, we explore the robustness of edge-based methods for identifying functional networks related to sustained attention performance in a controlled task and use these networks to predict attentional fluctuations in a different, naturalistic paradigm. This framework provides a case study for the validity of novel edge-based methods as predictive tools.

In the current study, we leverage edge-based GLMs to demonstrate that high-frequency fluctuations in sustained attention to visual and auditory information involve not only similar univariate brain activity, but also similar functional network reconfiguration. Importantly, we find that univariate activity and functional networks provide unique insights into neural mechanisms of sustained attention, demonstrating utility of the edge-based approach. Finally, we show that edge-based networks generalize between distinct task contexts. Results highlight the robustness of time-resolved functional networks for the study of sustained attention fluctuations across unique perceptual modalities and tasks.

## Methods

Data was collected at the MRI Research Center at the University of Chicago. Participants (N=60) completed at least one session of a two-session fMRI study. Functional MRI data were part of a large data collection effort and sample size was selected to provide high power for multiple planned analyses. Sessions were completed approximately one week apart (mean time between sessions=10.88 days, SD=9.87 days). During both sessions, participants viewed or listened to naturalistic narrative stimuli, performed a sustained attention task, and completed an annotated rest task in the fMRI scanner. Scan order for a single session was as follows: single-modality (auditory-only or visual-only) naturalistic stimulus, annotated rest, audio-visual naturalistic stimulus, annotated rest, sustained attention task. Annotated rest task data are not analyzed here. Functional MRI data from the sustained attention and annotated rest tasks have been reported previously using analyses independent of the current manuscript (Corriveau et al., 2025a; Ke et al., 2025).

Functional MRI data were collected on a 3T Philips Ingenia scanner. Voxels were 2.526mm x 2.526mm x 3mm. Volumes were collected using a multiband sequence with a repetition time of 1 second. Three volumes were removed from the start of each scan. All study procedures were approved by the Social and Behavioral Science Institutional Review Board at the University of Chicago.

Following the fMRI scan, participants completed a series of behavioral tasks. Participants watched or listened again to the naturalistic narrative stimuli presented during scanning and provided a continuous engagement rating on a sliding scale (described in detail below). After the narrative, participants also provided one value on the sliding scale rating how engaging they found the stimulus overall. Prior to this second presentation of narrative stimuli, participants also completed memory tasks for the naturalistic stimuli as well as images and sounds presented during the sustained attention task. Memory data and summary engagement scores are not analyzed here.

### Audio-visual continuous performance task

Sustained attention performance was tested in both sessions using a 10-minute audio-visual continuous performance task (avCPT; Corriveau et al., 2025b). During the avCPT, participants were presented a stream of trial-unique images and sounds presented simultaneously. Images were presented continuously for 1.2 seconds each whereas sounds were presented for 1 second with a 200 ms inter-trial interval to allow participants to distinguish individual sounds. Participants made a category-frequency judgement on either the images or the sounds such that both modalities served as the task-relevant modality across the two scanning sessions. Order was counterbalanced across participants. Participants were instructed to press a button when a stimulus from the task-relevant modality belonged to the frequent category (indoor/outdoor images, natural/manmade sounds) and withhold a button press to infrequent stimuli.

In rare cases, participants may have responded slowly to a frequent-category stimulus after the onset of the following trial. In this situation, it would appear that the participant made two key presses during the following trial when the first key press was intended as a response to the preceding trial. To account for this, we performed response reassignment if the following conditions were met: 1. A participant responded more than once during a frequent-category trial, 2. the first response was made within 100ms of the trial onset, and 3. no response was made to the previous frequent-category trial. In these cases, the first key press would be attributed to the preceding trial. Response reassignment was rare in both visual-relevant (mean reassignments = 0.548, SD=0.861) and auditory-relevant (mean reassignments=3.81, SD=3.46), affecting fewer than 0.8% of trials.

Sustained attention performance was behaviorally quantified in two ways. Accuracy of responses to infrequent stimuli served as momentary probes of sustained attentional state, with correctly withheld responses indicating high attention and failure to withhold a response indicating a sustained attentional lapse. Additionally, sustained attentional state was evaluated continuously using variance time course (VTC) which identifies periods of consistent, in-the-zone responding and erratic, out-of-the zone responding (Esterman et al., 2013). The VTC quantifies deviation response times, such that low VTC values indicate periods of stable responding and therefore better sustained attention, whereas high VTC values are indicative of worse sustained attention. The VTC was computed using correct responses to relevant, frequent-category trials. Response time courses were linearly detrended to account for speeding over time. Response times were z-scored and converted to absolute values. Finally, missing trial responses were linearly interpolated using responses from the two adjacent trials.

Logistic mixed-effect models were used to test whether variability in response times predicted successful performance on infrequent trials during the avCPT. Models included a fixed effect of preceding VTC, calculated as the average VTC during the three trials preceding an infrequent probe trial, as well as a subject-level random intercept. Models were fit for auditory-relevant and visual-relevant runs separately.

### Naturalistic narrative stimuli

Participants were presented four naturalistic narrative stimuli across both scan sessions— an auditory-only podcast, a silent movie, and two audio-visual movies. The podcast (“*Pie*,” 16 min 34 s) consisted of a baker describing the process of baking a berry pie. The silent movie (“*Croissant*,” 16 min 59 s) featured clips from a baker in the process of making strawberry and cream-filled croissants. One audio-visual movie (“*Cake*,” 15 min 40 s) featured the baking of a tiered chocolate cherry cake with narration. The final audio-visual stimulus (14 min 49 s) was a clip from the movie *North by Northwest* including dialogue between two main characters and a suspenseful airplane chase scene.

Following both scans, participants watched or listened to each stimulus a second time and performed an engagement rating task. Task instructions were to continuously indicate their subjective level of engagement using arrow keys to adjust the position of a slider bar on a scale from “Not engaging at all” to “Completely engaging.” Participants could indicate a large change in engagement by pressing the shift key and an arrow key simultaneously. The sliding scale contained 25 steps and the bar always initialized on step 13. Single key presses moved the bar one step and large jumps (shift+arrow key) moved the bar 5 steps.

### fMRI preprocessing

Functional MRI data were preprocessed using Analysis of Functional Neuroimages (AFNI version 19.0). The first 3 TRs of each functional run were excluded from analysis. Despiking, slice-time correction, and motion-correction was applied to the data. We regressed covariates of no interest from the data, including a 24-parameter head motion model (6 motion parameters, 6 temporal derivatives, and their squares) and mean signal from subject-specific eroded white matter and ventricle masks and the whole brain. Additional motion censoring was performed on avCPT runs: volumes in which more than 10% of voxels were outliers or the Euclidean distance of the head motion parameter derivatives were greater than 0.25 mm were removed. We then registered the functional images to participants’ skull-stripped MPRAGE anatomical images with linear transformation and then normalized to the Montreal Neurological Institute (MNI) space with nonlinear warping.

Functional data for all runs were parcellated into 400 cortical and 32 subcortical nodes using the Schaefer 400-node atlas (Schaefer et al., 2018) and the Melbourne Subcortical Atlas Scale II (Tian et al., 2020), respectively. This resulted in 432 time series for each run.

### Data exclusion

Functional MRI data from avCPT runs were excluded if in-scanner head motion exceeded any of the following criteria: maximum framewise displacement after censoring > 4mm, average framewise displacement after censoring > 0.15mm, or greater than 50% of volumes censored. Of the 57 participants who completed an auditory-relevant run of the avCPT, 10 were removed for head motion. For visual-relevant avCPT runs, fMRI data was available for 56 participants, 9 of whom were removed for head motion. Visual-relevant data was excluded from one additional participant who was missing data from 134 ROIs (∼31% of the brain). Additionally, we did not analyze fMRI data from participants who performed more than 2.5 standard deviations below mean performance (*A’*) across all auditory and visual runs of the avCPT task. One participant’s auditory-relevant run and one participant’s visual-relevant run were removed for low performance. This left a total of 46 participants with auditory-relevant and 45 participants with visual-relevant fMRI data. Of these, 37 participants are included in both analyses.

Naturalistic fMRI runs were used for model testing. Therefore, to maximize the number of possible testing datasets and because excessive motion could only impair model generalization, we did not perform exclusions for head motion during naturalistic runs. Neural data from one participant watching *North by Northwest* and one participant listening to *Pie* were excluded because data was missing >100 nodes. Final analyses of fMRI naturalistic data were performed on the following sample sizes: *Croissant* N=57, *Pie* N=55, *Cake* N=57, *North by Northwest* N=58.

We excluded behavioral engagement rating data from participants who reported a change in engagement less than once per minute on average, suggesting task noncompliance. Three engagement ratings from *Pie*, three from *Croissant*, four from *Cake*, and three from *North by Northwest* were excluded due to infrequent responding. Mean engagement time courses were calculated from the following sample sizes: 54 from *Pie*, 55 from *Croissant*, 50 from *Cake*, and 54 from *North by Northwest*.

### Univariate activity related to sustained attention

We conducted a general linear model analysis on BOLD activity to identify brain regions that track fluctuations in sustained attention. We fit two first-level models with sustained attention regressors of interest at the ROI level (Fortenbaugh et al., 2018; Jones et al., 2024). The first model included three attention-related regressors: incorrect presses to relevant, infrequent targets (lapses), correctly-withheld responses to relevant, infrequent targets (correct omissions), and incorrect failures to respond to relevant, frequent stimuli. Trial types were modelled with impulse regressors. Individual participants’ time series were fit using the autoregressive (AR) model in nilearn. Additional impulse regressors were included in the model for each censored time point. Regressors were convolved with the SPM HRF model as well as its temporal derivative. To identify ROIs in which activity differed between high attention and lapses, we computed the contrast: correct omissions – lapses.

The second model sought to identify activity related to continuous fluctuations of sustained attention. This model included a parametric regressor of interest, the VTC, which provides a continuous measure of sustained attentional state. Additionally, a boxcar regressor of constant duration was used to model individual trials and a boxcar regressor of variable duration, in which the duration reflected the linearly-detrended response time on each trial (Mumford et al., 2023), was included to account for activity that reflects response speed alone. Again, models were fit using nilearn’s AR model and impulse regressors were included for censored time points. Regressors were convolved with the SPM HRF model and its temporal derivative. To identify regions whose activity fluctuated with sustained attention, we computed the contrast VTC over baseline.

To correct for family-wise error rates, max-T thresholding was performed for univariate analyses. T-statistics were converted to F-statistics for permutation testing. Observed F-statistics were compared to null distributions generated by permuting F-statistics across ROIs 10,000 times. Significant ROIs were those whose observed statistics were more extreme than the null distribution, *p*<.05.

### Edge co-fluctuation networks related to sustained attention

Edge co-fluctuation time series quantify high-frequency changes in activity synchrony between pairs of brain regions (Faskowitz et al., 2020; Zamani-Esfahlani et al., 2020). Intuitively, edge time series “unravel” the Pearson correlation by quantifying two regions’ simultaneous deflection from their mean activity level. Computing the average of an edge time series returns the Pearson correlation between two regions’ activity time courses. For all pairs of brain regions, we calculated the edge time series as the dot product between z-scored BOLD activity time courses. The resulting vector quantifies the extent to which, at each time point, the regions were activating (or deactivating) similarly relative to their mean.

Importantly, we can fit general linear models to edge time series to identify pairs of brain regions whose co-fluctuations track sustained attention performance (Jones et al., 2024). We fit the two models described above to edge co-fluctuation time series to identify edge networks involved in successful vs. lapsing sustained attentional states, as well as continuous attentional fluctuations using the variance time course.

Significant edges were those whose F-statistics were more extreme (p<.05) than a random null, created by sign-flipping the true edge values 10,000 times. To control for type I errors, we identified edges that significantly tracked sustained attention performance across participants in both auditory-relevant and visual-relevant runs. This set of edges can be thought of as a modality-general edge network, as it tracked sustained attention regardless of modality in which stimuli were presented. Significance of overlap between runs was evaluated comparing the true number of edges that appeared in both runs to a null distribution created by randomly permuting the rows and columns of the significant edge matrices in both visual-relevant and auditory-relevant runs and counting the number of edges that were shared between runs, 10,000 times.

To evaluate the contribution of canonical networks to sustained attention, significant edges were grouped into 8 networks (7 canonical networks from Yeo et al., 2011 and 1 subcortical network from Tian et al., 2020). The significance of canonical network involvement was calculated by comparing the number of significant edges belonging to a canonical network to a null distribution calculated by permuting rows and columns of edge matrices, 10,000 times. Significant networks were those whose true edge membership was greater than the shuffled distribution, Bonferroni corrected for the number of within- and between-network comparisons made, *p*<(.05/36).

### Edge networks predicted from univariate activity

To examine the unique contribution of edge co-fluctuations to predictive power of sustained attention performance, over and above contributions of ROI activity alone, we compared edge networks that tracked attention fluctuations to those predicted by univariate activity. Univariate predictions were calculated using the following formula (from Jones et al., 2024):

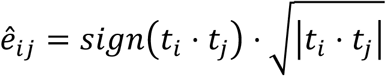

This calculation provides a theoretical edge map that should be expected based on the assumption that if activity in two regions is similarly related to a third variable of interest (in this case, sustained attention performance), their co-fluctuation time series should also be related to this third variable. If the edges predicted in this theoretical edge map largely overlap with the edges observed to track sustained attention performance, it would suggest that edge-based predictions are redundant with ROI-based predictions. However, if ROI activity-based predictions are largely unique from observed edge relationships, it would suggest that edges provide additional explanatory power above and beyond univariate activity.

### Generalization of edge networks to predict narrative engagement

To test whether edge networks defined during a controlled sustained attention task also track attentional fluctuations in more naturalistic contexts, we compared predicted changes in attentional state based on edge network strength to subjective reports of engagement while participants watched and listened to naturalistic narratives. Using a leave-one-subject-out approach, we identified modality-general edge networks—that is, edges that were related to sustained attention successes vs. lapses or VTC in both auditory- and visual-relevant task runs— for N-1 subjects. Then, we calculated TR-by-TR edge network strength in the left-out subject while they viewed and/or listened to each of the four naturalistic narrative stimuli. Edge network strength was defined as the difference in average edge co-fluctuation strength between positive (those with positive T-values) and negative (those with negative T-values) edges. This edge network strength calculation is based on previous work (Rosenberg et al., 2016) but is adapted to return the network strength at each time point. The resulting time course reflects time-varying network strength across an entire naturalistic run.

We then correlated the modality-general edge network strength time series with subjective ratings of engagement across an entire run. If edge networks generalize to capture subjective engagement, this correlation should be stronger than a null distribution, created by circle-shifting the true or reversed network strength time course 10,000 times. Circle shifting was limited such that the shift had to be greater than 8 time points from the true time course. For main analyses, edge network strength was calculated at the participant-level and compared to the mean engagement time course across all participants, as the group mean may reflect a less-noisy estimate of stimulus-driven engagement (Song et al., 2021). Mean engagement was computed by z-scoring engagement time courses within participants and averaging time courses across participants. We include results where each individual’s edge network strength time course was correlated with their unique z-scored subjective engagement time course in the supplement.

## Results

Performance on the avCPT, measured as *A’*, was high during both visual-relevant (M=0.937, SD=0.045) and auditory-relevant runs (M=0.744, SD=0.116), demonstrating participant compliance. A logistic regression predicting avCPT performance with a main effect of run type and a participant-level random intercept revealed that performance was significantly higher during visual-relevant runs (*brun type=*0.192, *SE=*0.019, *p<*.001) than auditory-relevant runs, suggesting increased difficulty in the auditory task. Similar results were observed in a larger behavioral sample (Corriveau et al., 2025b).

Response time variance, quantified using the VTC, significantly predicted success on infrequent trials during the avCPT. Specifically, infrequent probes in which participants correctly withheld a button press (correct omissions) were preceded by more stable responding, and therefore lower preceding VTC, in both visual-relevant (*b*=-0.141, SE=0.053, p=7.69*10^-3^) and auditory-relevant sessions (*b*=-0.135, SE=0.046, p=3.33*10^-3^). These findings confirm that the VTC tracks ongoing fluctuations in sustained attention performance, replicating previous work (Esterman et al., 2013).

### Auditory and visual sustained attention fluctuations recruit shared activity

To investigate univariate activity involved in sustained attention, we explored evoked activity for the following GLM regressors: correct omissions vs. lapses and the variance time course. Thresholded univariate maps are visualized in **Figure 1**. Unthresholded maps are included in **Supplementary Figure 1**.

**Figure 1.**
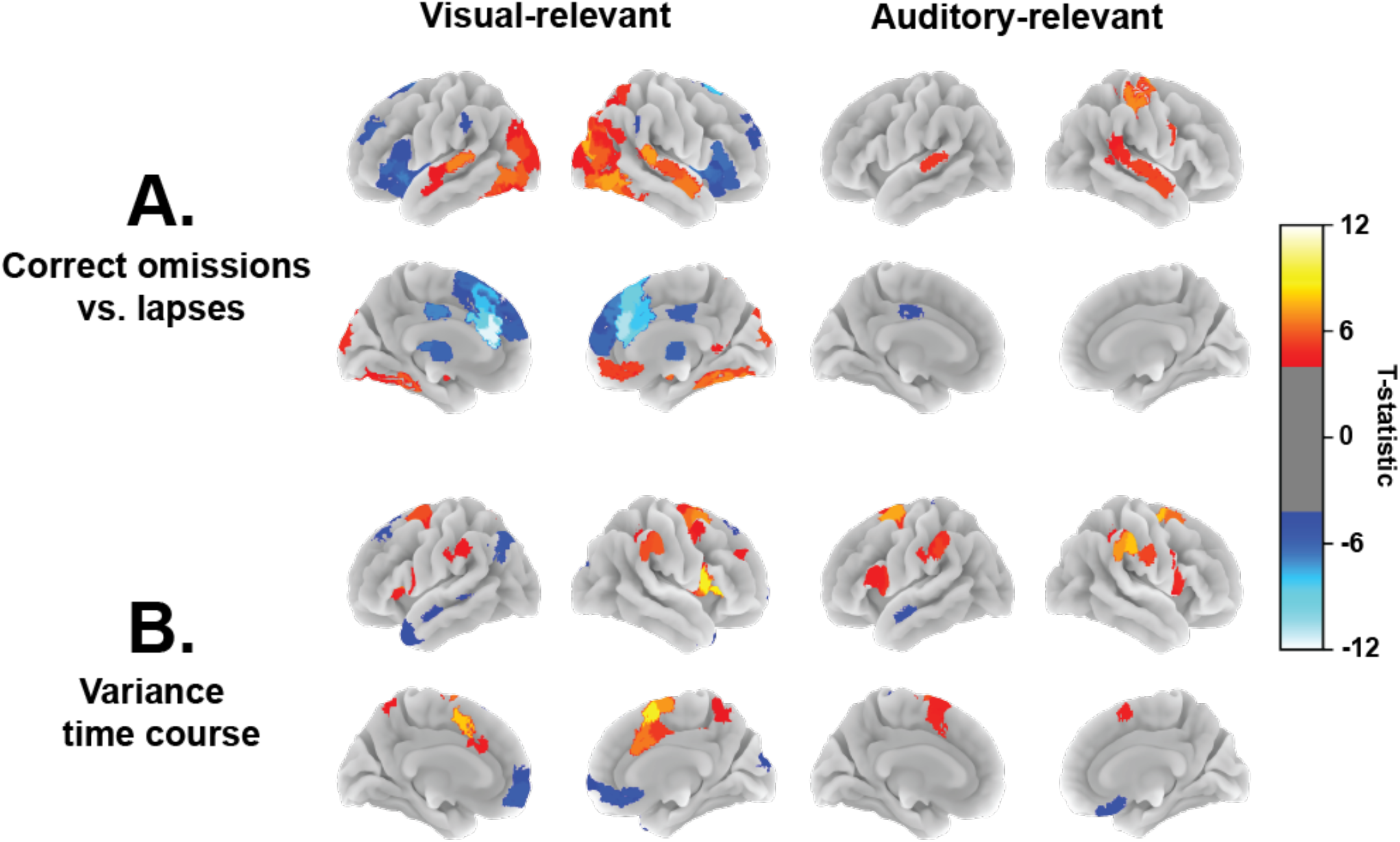
Maps visualizing significant results from contrasts of interest. (A) Difference maps for correct omissions vs lapses reveal regions in the temporal cortex and the PCC involved in both visual- and auditory-relevant sustained attention. Activity more strongly evoked by correct inhibition of responses to infrequent trials is visualized in warm colors, and activity related to failure to inhibit responses is visualized in cool colors. (B) Univariate activity associated with response time variance reveal regions in the dorsal and ventral attention networks that track sustained attention fluctuations. Regions in warm colors indicate areas whose activity increased with greater response time variance, reflecting worse sustained attentional states. Cool colors reflect regions whose activity increased with decreases in response variance, indicating better sustained attention. Colored regions reflect those surviving max-T thresholding, *p*<.05.

We first investigated activity differentially related to correct omissions vs. failures of sustained attention (lapses) on infrequent trials (**Figure 1A**). In visual-relevant runs, visual and temporal regions were involved in successful sustained attention performance, whereas areas in the prefrontal cortex, frontal operculum/insula, posterior cingulate cortex (PCC), and subcortex were more strongly activated during failures of sustained attention. This pattern of results largely replicates previous work (Fortenbaugh et al., 2018; Jones et al., 2024). During auditory-relevant runs, successful sustained attention performance was characterized by increased activity in temporal and right-hemisphere somatomotor regions, while increased left-hemisphere PCC activity was associated with lapses. Regions common to both tasks included bilateral temporal areas involved in sustained attentional successes and left PCC involved in lapses.

We also examined activity related to the variance time course (**Figure 1B**). Regions showing positive statistics reflect areas whose activity increased with greater response time variance, or worse sustained attentional states, while regions showing a negative relationship are those whose activity increased with reductions in response variance, indicating better sustained attentional states. In both visual-relevant and auditory-relevant tasks, regions in attention and salience networks, such as the frontal operculum/insula, temporal parietal junction (TPJ), frontal eye fields (FEF), and anterior cingulate cortex (ACC), showed increased activity under worse sustained attentional states. These results closely replicate previous work (Esterman 2013; Fortenbaugh et al., 2018; Jones et al., 2024).

### Shared edge networks underlie auditory and visual sustained attention

We next examined how network reconfiguration at the TR-by-TR time scale tracked changes in sustained attention. We fit our GLMs to edge co-fluctuation time series to identify edge networks related to our sustained attention contrasts of interest, i.e., correct omissions vs. lapses and the variance time course. Models were fit for visual-relevant and auditory-relevant fMRI runs separately. Resulting networks comprise the set of edges whose co-fluctuation time course significantly reflected changes in sustained attention during the avCPT.

Contrasting correct omissions vs. lapses revealed a set of 5,274 edges whose co-fluctuation was stronger during correct omissions and 6,654 edges whose co-fluctuation was stronger during attentional lapses during visual-relevant runs (**Figure 2A**). In auditory-relevant runs, 1,883 edges showed stronger co-fluctuation during correct omissions and 2,739 edges showed stronger co-fluctuation during lapses. Between visual-relevant and auditory-relevant runs, 160 edges positively tracking correct omissions and 271 edges positively tracking lapses were shared by both run types, a significant amount of overlap in both cases (both *p*s<.001; **Figure 2B**). In this modality-general overlap, edges connecting the ventral attention to default mode and limbic networks showed higher co-fluctuations during correct omissions. Conversely, edges within the default mode and ventral attention networks, as well as edges between the somatomotor network and subcortex showed stronger co-fluctuations during attentional lapses. Edge networks also shared edges in the unexpected direction, although, as predicted, this overlap was not significant. The network that positively tracked correct omissions in visual-relevant runs shared 91 edges with the network that tracked lapses during auditory-relevant runs, whereas the network that tracked correct omissions during auditory-relevant runs shared 69 edges with the visual-relevant lapse network (*p*s>.999).

**Figure 2.**
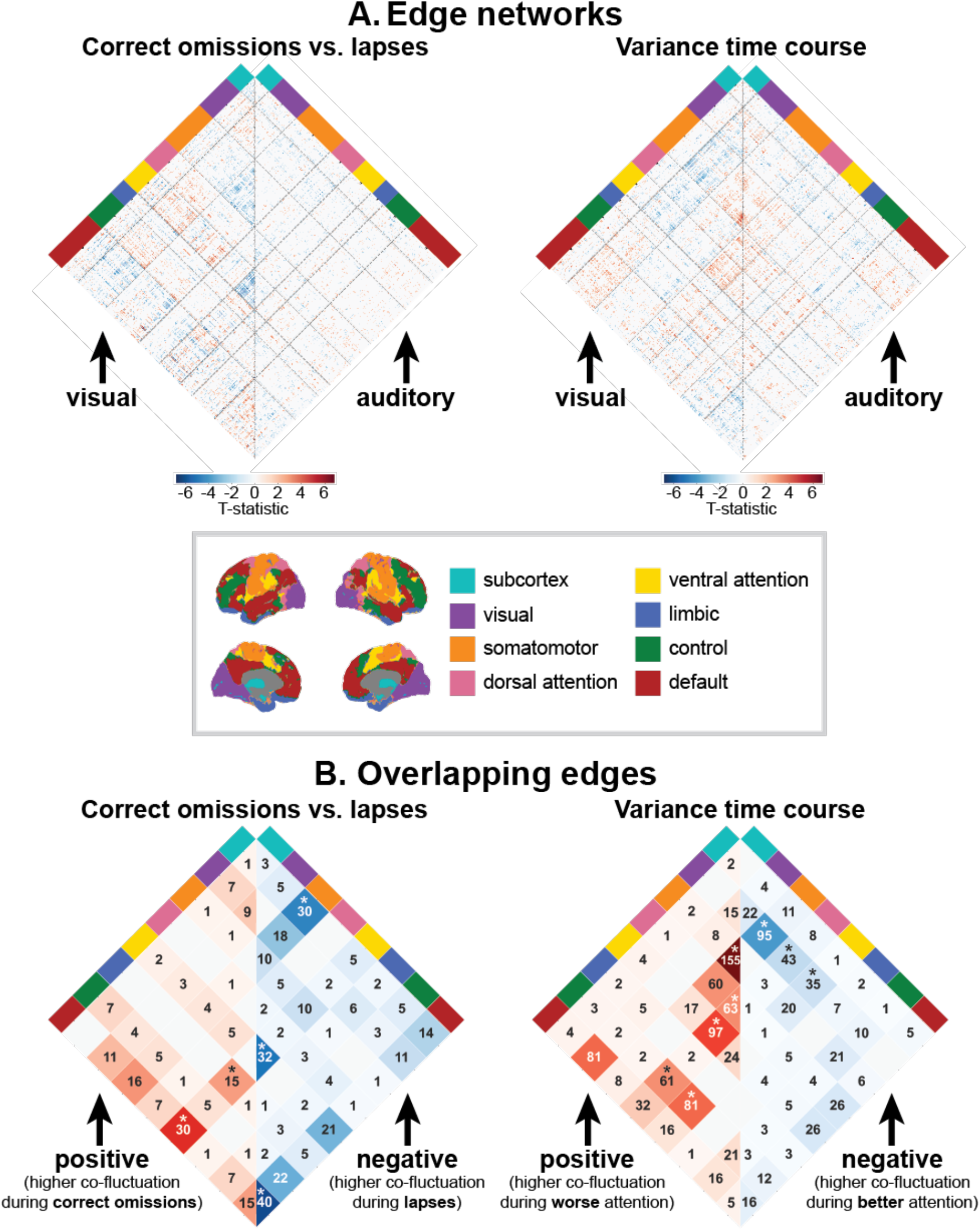
(A) Distributed edge networks are involved in sustained attention to visual and auditory tasks. Colored cells reflect pairs of regions whose co-activation time course was significantly positively (red) or negatively (blue) related to the contrast of interest. Note that only significant edges are visualized. (B) Overlapping edges were involved in sustained attention during both visual-relevant and auditory-relevant runs. Edge networks are summarized using canonical cortical and subcortical networks (Yeo et al., 2011; Tian et al., 2020). Stars (*) in B indicate networks with greater edge involvement than expected by chance.

When contrasting the variance time course against baseline in visual-relevant runs, 5,306 edges were positively related to the variance time course, i.e., showed higher co-fluctuation during worse sustained attentional states, whereas 5,507 edges showed a negative relationship with the variance time course. In auditory-relevant runs, 3,675 edges were positively related to the variance time course while 3,356 were negatively related. Both positive and negative edge networks showed significant overlap between the two sessions (790 overlapping positive edges, 403 overlapping negative edges; *p*s<.001). Overlapping edges positively associated with the variance time course, therefore showing stronger co-fluctuation during worse sustained attention, were found within the somatomotor and dorsal attention networks, between the dorsal attention and ventral attention networks, and between control networks and dorsal and ventral attention networks. Edges connecting the visual network to somatomotor, dorsal attention, and limbic networks were negatively associated with the variance time course, i.e., showed higher co-fluctuations during better sustained attention. Again, networks trained to differentially predict the variance time course between modalities did not significantly overlap. The network that positively tracked variance time course fluctuations during visual-relevant runs shared 58 edges with the negative network during auditory-relevant runs, while the negative network during visual-relevant runs shared 57 edges with the positive network during auditory-relevant runs (*p*s>.999). Results confirm that edge-based GLMs identify modality-general networks that reconfigure with fluctuations in sustained attention.

Interestingly, edge networks related to better sustained attention, that is, positive networks related to correct omissions vs. lapses and negative networks related to the variance time course, significantly overlapped in visual-relevant runs (1450 overlapping edges, *p*<.001) but not auditory-relevant runs (84 overlapping edges, *p*>.999). Similarly, networks related to worse sustained attention as measured by correct omissions vs. lapses and the variance time course were significantly overlapping in visual-relevant runs (1932 overlapping edges, *p*<.001) but not auditory-relevant runs (196 overlapping edges, *p*=.835). Edges related to sustained attention across all networks were not significant (0 edges related to better sustained attention, 3 edges related to worse sustained attention, *p*s>.999), likely due to limited overlap between auditory-relevant runs.

### Edge networks are not predicted by univariate activity

Does an edge-based GLM approach offer new insight into neural mechanisms of sustained attention fluctuations? Or, are the co-fluctuations observed to track sustained attention simply byproducts of activity in regions previously reported to underscore attentional fluctuations? To test these questions, we compared networks obtained from our edge-based GLM analysis to a network of edges predicted by univariate activity, by calculating the signed square root of univariate T-statistics between all pairs of ROIs. This calculation provides a theoretical edge network based on the assumption that, if activity in two ROIs varies as a function of sustained attention, their co-fluctuation should also track sustained attention.

Here, we compare theoretical modality-general edge networks—i.e., edges predicted to be related to sustained attention in both visual and auditory sessions—to observed networks. Theoretical edge maps based on univariate activity predicted large networks of co-fluctuation involved in sustained attention. Despite this, these predicted networks were largely independent of edge networks observed to underlie sustained attentional fluctuations (i.e., the networks in **Figure 2B**). Univariate activity-predicted networks included 4,454 edges stronger during correct omissions and 1,030 edges stronger during lapses across modalities. However, none of these edges were shared with the networks identified by edge-based GLM analyses (160 edges involved in correct omissions, 271 edges involved in lapses), indicating that theoretical edge predictions missed 100% of observed edges. When contrasting the variance time course against baseline, univariate activity predicted networks of 5,877 edges positively and 1,978 edges negatively related to the variance time course. Of these, 417 positive edges and 33 negative edges were also identified by edge-based GLMs, suggesting that theoretical edge predictions missed 47.2% and 91.8% of observed edges, respectively. In combination, results suggest that edge-based GLMs identify unique neural substrates not predicted by univariate activity alone.

### Edges related to sustained attention lapses also track subjective engagement

Lastly, we tested whether the modality-general networks involved in sustained attention during a controlled task generalized to predict engagement during naturalistic narrative stimuli. Narrative engagement, while related to sustained attention, was measured subjectively and continuously by participants themselves as they watched or listened to movies and a podcast. Successful generalization of edge networks requires that they capture features of attentional fluctuations that are not task-dependent.

Using a leave-one-subject-out approach, we calculated the network strength time course in modality-general edges while participants viewed one silent movie (*Croissant*), one podcast (*Pie*), and two audio-visual movies (*Cake* and *North by Northwest*) in the fMRI scanner. We compared this predicted neural time course to the mean time course of subjective engagement provided by participants using Pearson’s correlation. Significance was determined by comparing the average observed prediction (*r-*values) to a null distribution created by circle-shifting the predicted edge time courses within subjects 10,000 times. Mean engagement time courses are visualized in **Supplementary Figure 2**.

Edges related to correct omissions vs. lapses during auditory and visual runs of the avCPT also predicted fluctuations in subjective engagement during all narratives (*p*s<.05; **Figure 3A**). As expected, network strength time courses were positively correlated with ratings of subjective engagement across participants, suggesting that co-fluctuation related to better attentional states during the controlled avCPT was also stronger during moments of increased subjective engagement. Correlations between network strength and subject-level engagement time courses were less reliable, but still significant in both audio-visual movies (**Supplementary Figure 3A**).

**Figure 3.**
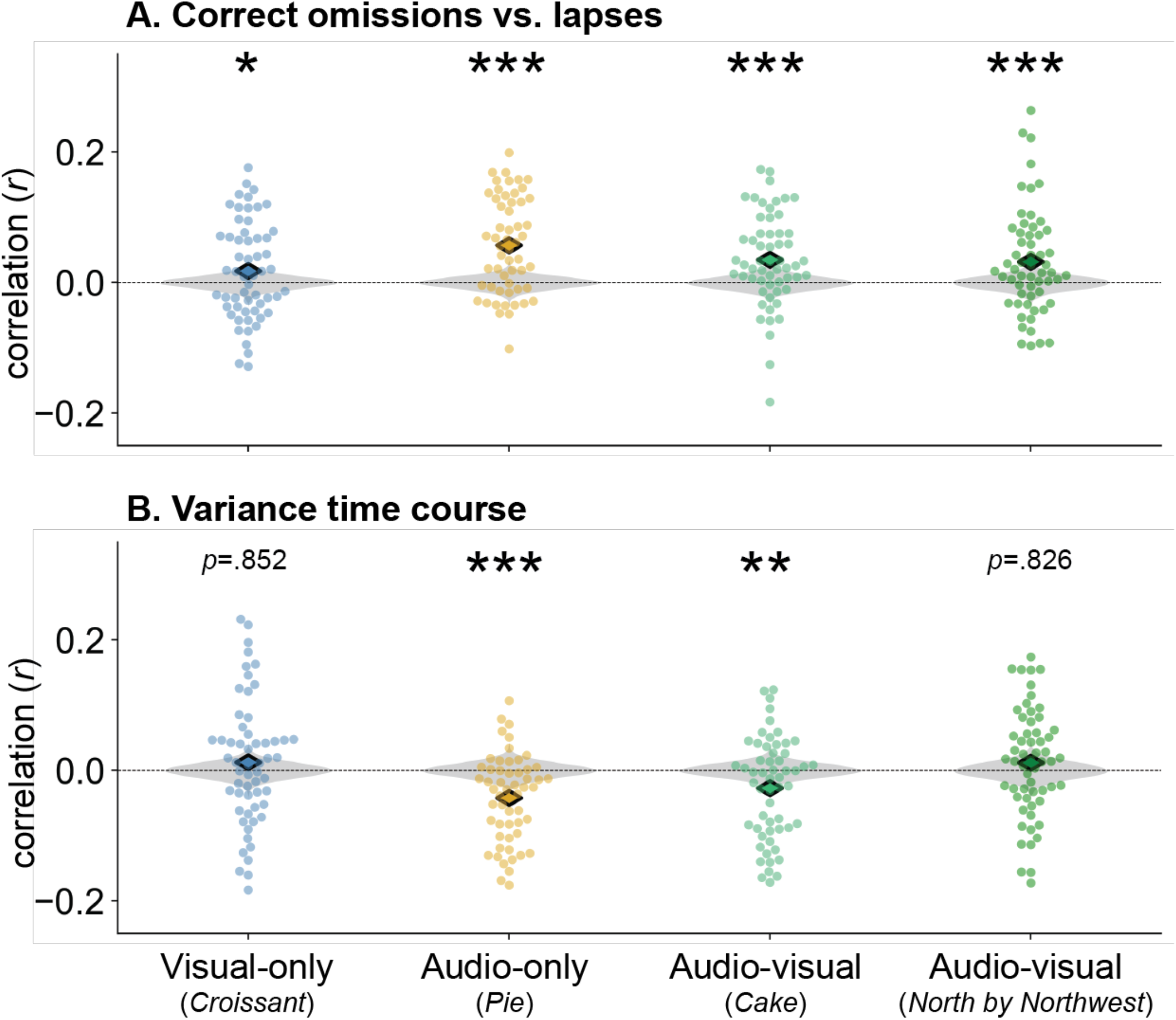
Modality general networks related to (A) correct omissions vs. lapses and (B) variance time course fluctuations during the controlled avCPT also captured fluctuations in subjective engagement. Dots reflect individuals’ correlation between network strength and mean engagement time courses. Colored diamonds reflect the observed mean correlation across participants. Gray null distributions were created by circle-shifting network strength time courses within participants and recalculating the mean correlation 10,000 times. *p<.05, **p<.01, ***p<.001.

Modality-general edges related to the variance time course predicted subjective engagement during the podcast (*p*<.001) and one audio-visual movie (*Cake*; *p*=4.40*10^-3^), but not the silent movie or audio-visual *North by Northwest* (**Figure 3B**). The correlation between subjective engagement and VTC edge network strength time courses was predicted to be negative, because edges in the VTC network are inversely related to sustained attention. When predicting subject-level engagement time courses, correlations were less strong but remained significant in the audio-visual *Cake* (**Supplementary Figure 3B**). In combination, results show that edges related to time-resolved measures of sustained attention during a controlled task also capture fluctuations in attentional engagement in a less-controlled, naturalistic paradigm.

## Discussion

The current study examined the ability of time-resolved edge co-fluctuations to capture dynamic fluctuations of sustained attention from moment-to-moment. Univariate and edge-based GLMs identified a set of core, modality-general regions and networks related to sustained attention regardless of the perceptual modality of the task. Further demonstrating the robustness of edge-based models, modality-general edges also predicted moment-to-moment changes in subjective engagement while participants watched movies and listened to a podcast. Results support the utility of edge time series in the identification of brain networks involved in dynamic attention, cognition, and behavior.

Univariate activity involved in sustained attention largely replicated findings of sensory and attention region involvement from previous work (Fortenbaugh et al., 2018; Jones et al., 2024; Eickhoff & Langer, 2013). Interestingly, the use of an audio-visual CPT in the current study revealed that, during visual-relevant sessions, activity in both visual and auditory regions was increased when participants were performing well, despite auditory stimuli being irrelevant in these sessions. While the interpretation of increased activity is not straightforward, these results are in line with behavioral work finding that processing of irrelevant stimuli (in addition to relevant stimuli) increases under better sustained attentional states during CPTs (Corriveau & Chao et al., 2024; Corriveau et al., 2025b). Additional work is required to examine how processing may be changing at a neural level.

Using edge-based GLMs, we identified modality-general networks whose co-fluctuation tracked trial-to-trial fluctuations in sustained attention. Edge networks were reliable across perceptual modality, mirroring previous work suggesting that networks that predict individual differences in sustained attention contain core modality-agnostic components (Corriveau et al., 2025a) and demonstrating robustness of edge-based GLM results. Further validating edge-based GLM methodology, networks identified using this time-resolved approach replicate findings from previous methods using functional connectivity averaged over longer time periods. For example, previous work has associated worse sustained attentional states with increased connectivity in the default mode network (Kucyi et al., 2017), in line with our current findings. Additional work using static functional connectivity suggests that increased connectivity within the somatomotor network predicts worse sustained attention across individuals (Corriveau et al., 2025a; Lyu et al., 2025; O’Halloran et al., 2018), a finding we also observed in the current study. This suggests that edge networks, which were defined on temporally-resolved and therefore potentially much noisier patterns of neural activity, capture robust neural substrates underlying sustained attention.

Fewer regions and edges were identified when contrasting correct omissions vs. lapses in auditory-relevant runs relative to visual-relevant runs. While this may reflect key differences between neural activity between perceptual modalities, this may also be the result of worse behavioral performance and possibly greater difficulty during auditory-relevant runs. Errors during auditory-relevant runs may not only reflect lapses in sustained attention but also a failure to correctly categorize a trial. This possibility is also supported by the observation that the edges related to correct omissions vs. lapses were unique from those related to the variance time course in auditory-relevant runs. Edges in the network tracking correct omissions vs. lapses may reflect not only sustained attention but also processes required by the increased difficulty such as effort, which may not be reflected in the variance time course network. Future work may seek to balance task difficulty across modalities to disentangle these possibilities.

Modality general edge networks also generalized to capture fluctuations in engagement during naturalistic narratives, despite many differences between the controlled avCPT and naturalistic task contexts. Viewing and listening to naturalistic narratives in the scanner was passive, such that participants watched or listened to the stimuli with no experimenter-imposed task goals. While related to sustained attention, narrative engagement may reflect a confluence of phenomena, including participants’ internal states and the emotionality of stimuli (Song et al., 2021). Further, narrative engagement was reported subjectively, requiring that participants have accurate insight into their own cognition. Even with these limitations, edge networks trained to predict objective measures of sustained attention, in particular correct vs. incorrect responding, also predicted fluctuations in engagement. Previous work using dynamic sliding windows found that a network trained on static functional connections related to sustained attention predicted ongoing engagement during an audio-visual movie but not an auditory-only podcast (Song et al., 2021). The time-resolved approach method used in the current analyses therefore may strengthen the robustness and generalizability of functional networks across task contexts.

The current results demonstrate the utility of edge-based analyses for the study of neural mechanisms underlying dynamic mental states. Indeed, edge time series are ideal for investigating highly dynamic aspects of attention, cognition, and behavior that may fluctuate at different time scales. Importantly, the edge networks identified using GLMs were not the same as those predicted by univariate activity, despite the fact that edge co-fluctuation time series are mathematically equivalent to the interaction between univariate time courses (Merritt et al., 2024). This suggests that the extent to which the joint fluctuation of two regions tracks sustained attention is unique from those regions’ individual relationships with sustained attention. Therefore, examining both univariate responses and co-fluctuations using a GLM framework allows us to link activity and networks related to phenotypes of interest while still quantifying unique contributions from the two (Jones et al., 2024). This useful tool has immense potential for furthering understanding of dynamic attention and cognition.

## Conclusion

Univariate activity and edge-based GLMs identified brain regions and networks related to moment-to-moment fluctuations in sustained attention. Predictive networks were composed of shared edges regardless of the perceptual modality of the sustained attention task. Further, these modality-general networks were unique from those predicted by univariate activity and captured fluctuations in subjective attentional engagement in a vastly different task context. Together, this work demonstrates the utility of univariate and edge-based methods for the identification of unique neural substrates underlying dynamic sustained attention.

## Data and Code Availability

Data and code needed to replicate analyses are available at https://osf.io/xk6v9/

## Acknowledgements

The authors thank Henry M. Jones for providing open-source code used as the basis for main analyses and for thoughtful conversations on the current results.

## Author Contributions

CRediT: A.C.: Conceptualization, Formal Analysis, Visualization, Writing - original draft; J.K.: Data curation, Investigation, Writing - review & editing; M.D.R.: Conceptualization, Funding acquisition, Project administration, Resources, Supervision, Writing - review & editing

