## Supplementary materials for "Common brain network dynamics capture attention fluctuations in tasks and movies"

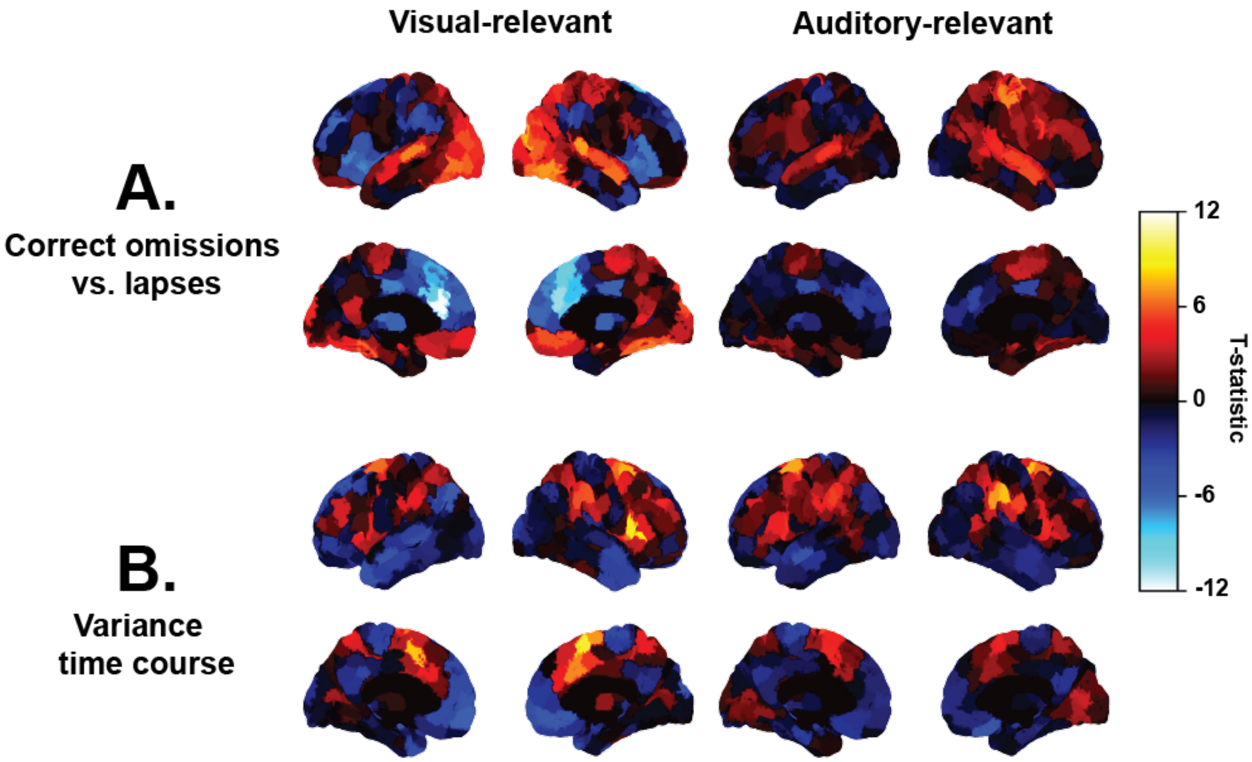


*Supplementary Figure 1.* Maps visualizing unthresholded results from contrasts of interest. (A) Activity more strongly evoked by correct inhibition of responses to infrequent trials is visualized in warm colors, whereas activity related to sustained attentional lapses is visualized in cool colors. (B) Univariate results from contrasting the variance time course against baseline. Warm colors indicate areas with increased activity related to greater response time variance, reflecting worse sustained attentional states. Cool colors reflect regions whose activity increased with decreases in response variance, indicating better sustained attention.

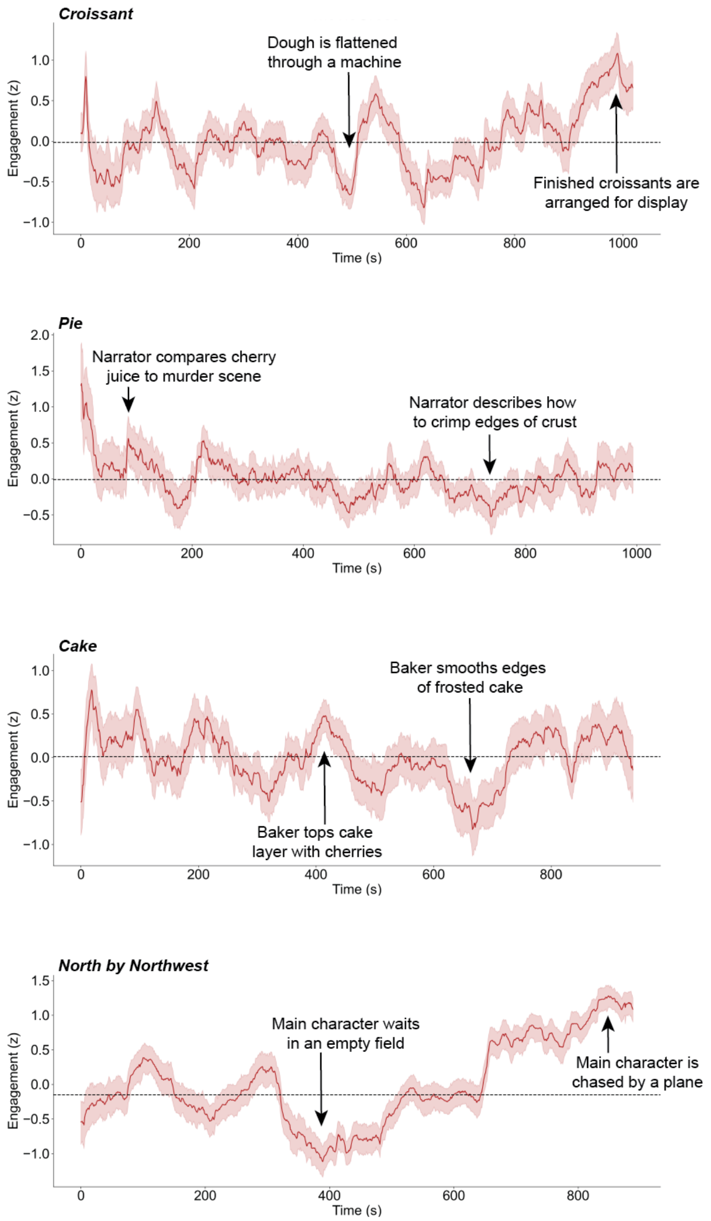


*Supplementary Figure 2.* Mean engagement time courses for naturalistic narrative stimuli averaged across participants. Time courses were z-scored within participants before averaging. Shaded regions reflect 95% confidence intervals. Dotted black lines indicate median engagement across the time course.

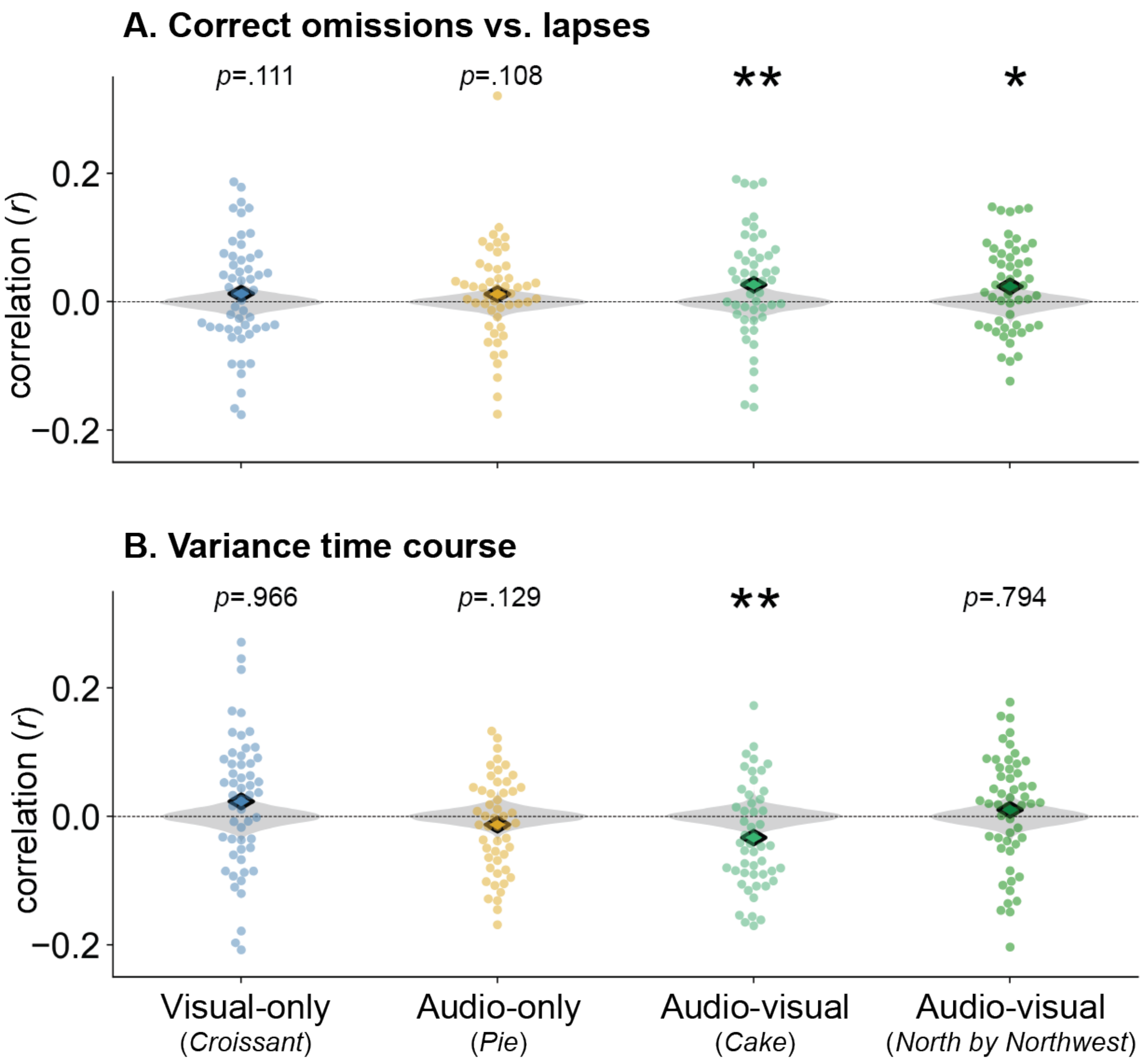


*Supplementary Figure 3.* Correlations between subject-level engagement time courses and predicted network strength time courses. Null distributions generated with circle-shifting are shown in gray. (A) Predictions generated from modality-general edges related to correct omissions vs. lapses were correlated with subject-level engagement in both audio-visual movies. (B) Modality-general edges related to the variance time course predicted subject-level engagement in one audio-visual movie, *Cake*.
